# A simplified co-culture reveals altered cardiotoxic responses to doxorubicin in hPSC-derived cardiomyocytes in the presence of endothelial cells

**DOI:** 10.1101/2025.02.21.639078

**Authors:** Marcella Brescia, Andrea Chatrian, Paul Keselman, James Gallant, Elsa Sörman Paulsson, Mervyn P.H. Mol, Rickard Sjögren, Karine Raymond, Valeria Orlova, Kalpana Barnes, Richard Wales, Jonas Austerjost, Michael W. Olszowy, Christine L. Mummery, Berend J. van Meer, Richard P. Davis

## Abstract

Cardiotoxicity is a significant challenge in cancer therapies, particularly with doxorubicin, a widely used anthracycline known for its broad anti-cancer spectrum but life-threatening cardiac side effects. There is a critical need for more predictive *in vitro* models to understand doxorubicin-induced cardiotoxicity and patient-specific drug responses. In this study, we used human pluripotent stem cell (hPSC)-derived cardiomyocytes (hPSC-CMs), cardiac fibroblasts (hPSC-cFBs) and endothelial cells (hPSC-ECs) to investigate the cardiotoxic effects of doxorubicin in two-dimensional mono-and multi-cell type cultures. By mimicking the cumulative dose effect seen in patients through repeated doxorubicin treatments and using a machine learning-based *in silico* image analysis tool, we could precisely quantify caspase 3/7 activity as an early toxicity marker and identify hPSC-CMs in multi-cell type cultures. This innovative approach allowed continuous monitoring of apoptosis from phase-contrast images, revealing that hPSC-ECs showed higher sensitivity to doxorubicin than isogenic hPSC-CMs or hPSC-cFBs and significantly enhanced cardiomyocyte toxicity in co-culture. In contrast, dermal fibroblasts differentiated from the same hPSC line showed no toxic response under the same treatment regimen. These results challenge the conventional focus on cardiomyocytes as the target of drug-induced cardiac damage. Our findings not only highlight the complex interplay among different cardiac cell types in mediating the toxic effects of doxorubicin, but also demonstrate the potential of AI-enabled tools to advance personalized drug screening and safety assessments.

## Introduction

Predicting patient-specific drug responses and adverse reactions remains a significant challenge in healthcare, particularly in the context of cardiotoxicity. This is most notable with chemotherapeutics, which can have well-known, life-threatening side effects with late heart failure evident in up to 10% of patients [1]. The variability in drug efficacy and safety, even among patients with identical diagnoses, complicates the prediction of such adverse effects [2]. The heart is one of several organs for which regulatory authorities mandate comprehensive cardiotoxicity assessments throughout the drug development process [3]. Conventionally, these assays have predominantly relied on *ex vivo* assays and *in vivo* studies in animals. However, the physiological and anatomical differences between humans and most animal models often limit the translatability of such studies, contributing to high failure rates (∼90%) of drug candidates in clinical trials due to unforeseen toxicities and lack of efficacy [4,5]. Contemporary regulatory initiatives, exemplified by the FDA Modernization Act 2.0, are driving a paradigm shift away from reliance on animal models towards the adoption of cell-based assays and advanced data analysis methodologies [6]. Human pluripotent stem cells (hPSCs) are emerging as a promising alternative, potentially offering more predictive models for evaluating drug-induced cardiotoxic effects and reducing reliance on animals [7]. Moreover, these cells are particularly valuable for personalized screening and improving understanding of why individual patients respond differently to cardiotoxic compounds [8].

Doxorubicin (Doxo), a widely used anthracycline, exemplifies the delicate balance between therapeutic efficacy and cardiotoxic risk. Despite its proven effectiveness in treating a variety of cancers, including acute leukemia, lymphomas, and various solid tumors in both adults and children, it is also associated with significant adverse effects, most notably cardiotoxicity [9]. This cardiotoxic effect, characterized by a spectrum of cardiac dysfunctions such as congestive heart failure, arrhythmias, and reduced left ventricular ejection fraction [10], manifests in a cumulative and dose-dependent manner and severely limits its repeated or long-term clinical use. The mechanisms involved are complex and not fully understood, but include mitochondrial dysfunction, calcium overload, DNA damage, and apoptosis following caspase activation [11-13].

hPSC-derived cardiomyocytes (hPSC-CMs) have become a mainstream tool for cardiotoxicity testing by both the pharmaceutical industry and academia, owing to their scalability and suitability for high-throughput safety assessments [14,15]. However, despite their widespread adoption, these models still present limitations in accurately predicting clinical outcomes. Increasing their complexity by incorporating additional hPSC-derived cardiac cell types such as endothelial cells (ECs) and cardiac fibroblasts (cFBs), has been shown to improve their physiological relevance and, consequently their clinical translatability [14]. This improvement is particularly evident in 3D cultures, for example microtissues or engineered heart tissues (EHTs), which not only promote the maturation of hPSC-CMs but also the predictiveness of drug responses [7,16], including the effects of Doxo on contraction dynamics [2,17].

However, 3D models are relatively inaccessible, especially to investigate cell-specific response in real-time. In this context, we investigated the potential of 2D multi-cell type cultures to offer a more predictive model than 2D hPSC-CM monocultures, while remaining more accessible than 3D cardiac models for both drug exposure and imaging. To analyze the dynamics of cardiotoxicity over prolonged and repeated exposures to Doxo and quantitatively compare the effects between different cell types, we applied a machine learning (ML)-based analysis tool to assess caspase 3/7 activity as a marker of early toxicity. This *in silico* tool could also identify hPSC-CMs in multi-cell type cultures, allowing us to track cardiomyocyte apoptosis in different cellular environments over time. Our findings revealed that the presence of specific cell types, notably hPSC-ECs, influenced the sensitivity of hPSC-CMs to Doxo and may be a key driver of cardiomyocyte toxicity. This study not only sheds light on Doxo-induced cardiotoxic mechanisms but also underscores the potential of AI tools to advance high-throughput analysis and personalized drug screening, paving the way for safer therapeutic interventions.

## Results

### hPSC-CMs co-cultured with other cardiac cell types appear more sensitive to Doxo

To emulate the *in vivo* pharmacodynamics of Doxo [18,19], an *in vitro* treatment protocol was used in which the hPSC-derived cells were exposed to Doxo for 4 h intervals every 48 h (Fig. 1A). Because the free diffusible Doxo concentration *in vivo* varies between 20 nM and 2 μM [20], we examined a range of concentrations (0.01 – 10 μM), with representative phase contrast images and analysis presented in Supplementary Fig. 1. We selected 1 μM Doxo as a proxy for the cardiotoxic effect seen *in vivo* since this concentration induced cumulative toxicity without immediate cell death in hiPSC-CMs.

**Figure 1.**
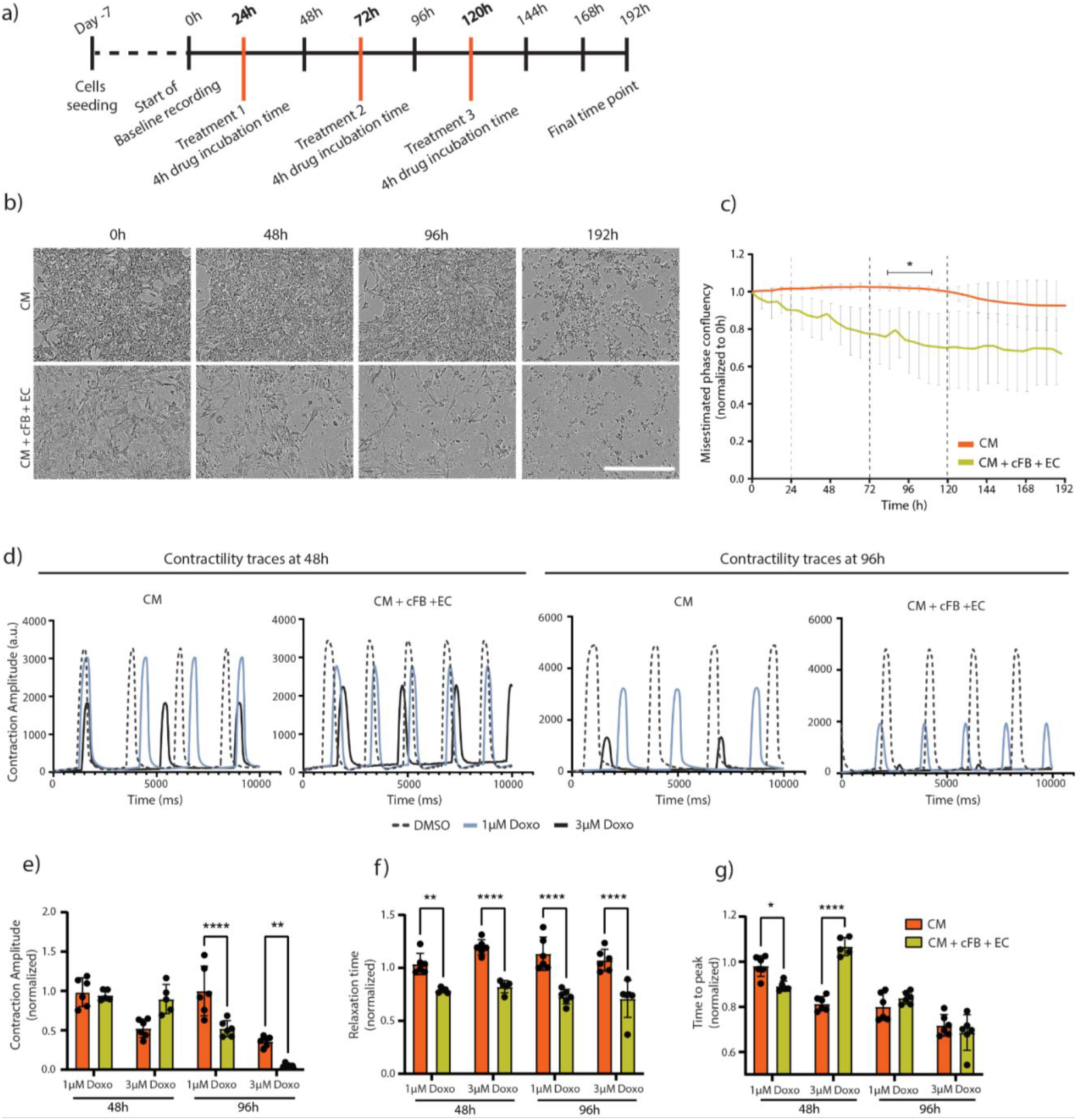
Cumulative Doxo treatment effect on hPSC-derived cardiac cell cultures. **a)** Schematic of treatment protocol timeline, with orange lines marking the time points of 4 h Doxo application to the cells. **b)** Representative phase contrast images of monoculture hiPSC-CMs (top panel) or co-cultured with hiPSC-cFBs and -ECs (bottom panel) treated with 1 µM Doxo. Images were acquired at time points corresponding to baseline (0h), 24 h after Treatment 1 and 2 (48h and 96h, respectively), and the final time point (192h). Scale bar, 200 μm. **c)** Quantification of phase confluency area calculated from live cell imaging, and normalized to baseline (0h), for conditions shown in **(b)**, highlighting limitations in accurately determining confluency. Dotted lines indicate treatment time points with 1 µM Doxo. The asterisk and black bar indicate the time points where there were statistically significant differences between the monoculture and the multi-cell type culture setups. **d)** Representative contraction traces of hiPSC-CMs in both monoculture and multi-cell type conditions at 48h and 96h (24 h after Treatment 1 and 2, respectively). Treatments were either DMSO (vehicle control), 1 µM or 3 μM Doxo. **e-g)** Graphs comparing contraction amplitude **(e)**, relaxation time **(f)**, and time to peak **(g)** of hiPSC-CMs in either monoculture or multi-cell type culture conditions at the indicated time points. Each was normalized to their respective vehicle control. Statistical significance was determined by two-way ANOVA analysis with Sidak’s multiple comparison test. Analysis is based on 3 biological replicates, each with 3 technical replicates, with error bars representing SEM.

When hiPSC-CMs were treated in co-culture with hiPSC-cFBs and hiPSC-ECs, Doxo induced toxicity more rapidly than in monocultures of hiPSC-CMs. In phase contrast images, cell death was immediately evident after a single 1 μM Doxo treatment in triple cultures, while monocultures of hiPSC-CMs required 3 cycles for visible cell death (Fig. 1B). Through frequent imaging, morphological changes indicating of apoptosis became evident, including cytoplasmic shrinkage, cell shape changes and cell detachment. However, when attempting to quantify the extent of toxicity, conventional phase microscopy was unable to distinguish live from dead cells (Supplementary Fig. 2A) due to cellular debris and clumping which affected the ability to apply size-or eccentricity-based threshold filters. This resulted in discrepancies with, for example, confluency measurements indicating only 20% loss in the co-culture condition at 96 h (Fig. 1C), despite most cells appearing dead (Fig. 1B). Nevertheless, we could distinguish differences in the toxicity dynamics between triple cultures and hiPSC-CM monocultures, with triple cultures exhibiting a more rapid onset of cell death following Doxo treatment (Fig. 1C).

**Figure 2.**
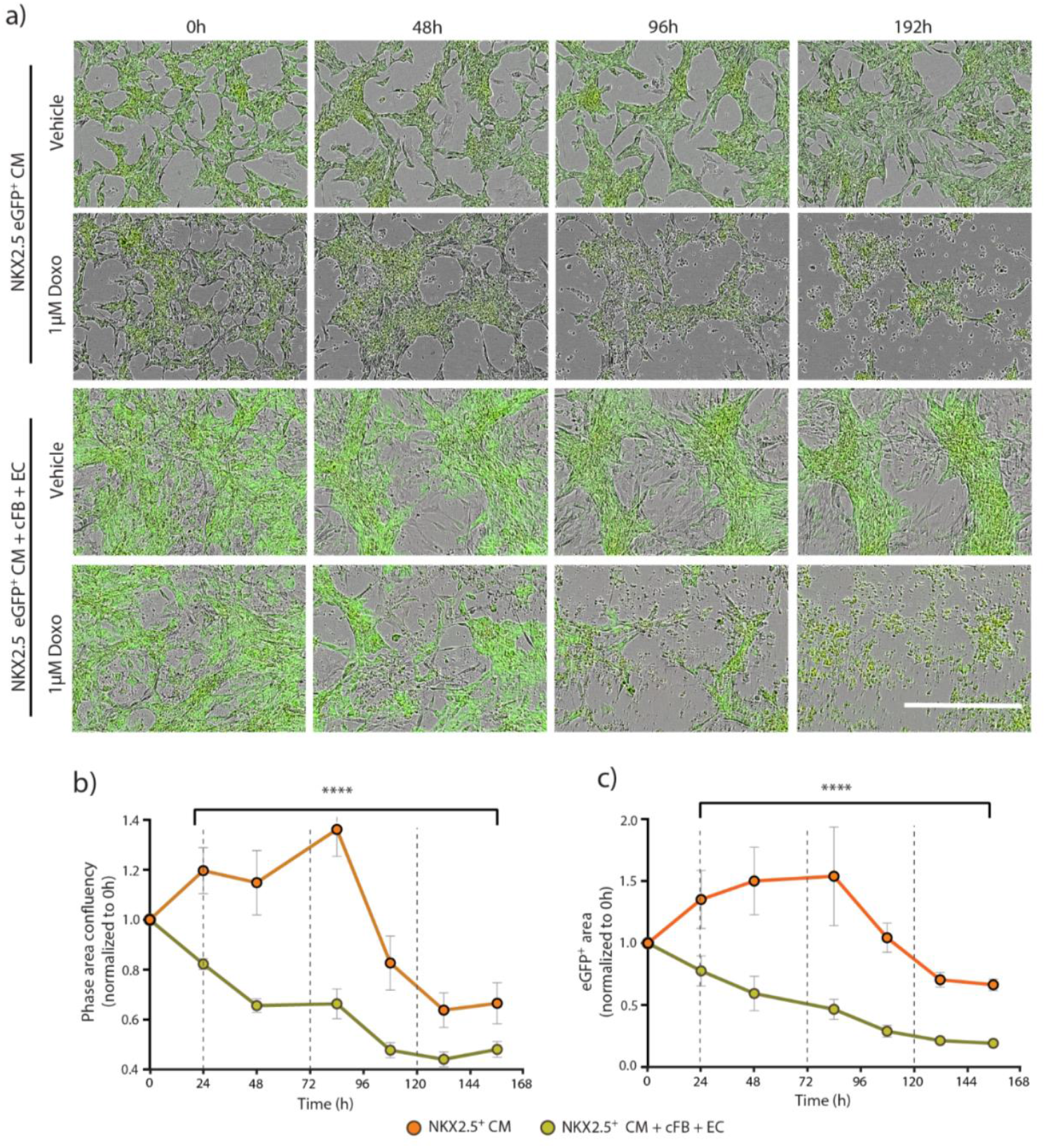
Cumulative Doxo treatment effect on hESC-CMs in mono-or tri-cellular cultures. **a)** Representative merged phase and green fluorescence images of NKX2.5-eGFP^+^ hESC-CMs in monoculture or co-cultured with hiPSC-cFBs and -ECs, treated with either 1 µM Doxo or Vehicle control (DMSO) following the cumulative treatment protocol. Images were acquired at time points corresponding to baseline (0h), 24 h after Treatment 1 and 2 (48h and 96h, respectively), and the final time point (192h). Scale bar, 200 μm. **b-c)** Quantification of phase **(b)** and green (NKX2.5^+^ hESC-CMs **(c)** confluency areas calculated from live cell imaging, and normalized to baseline (0h), for cell culture conditions outlined in **(a)** treated with 1 µM Doxo. Dotted lines indicate the treatment time points. Statistical analysis between cell culture conditions employed two-way ANOVA analysis with Sidak’s multiple comparison test for each time point. Analysis is based on 3 biological replicates, each with 3 technical replicates, with error bars representing SEM.

From video recordings of the hiPSC-cardiac cultures acquired during their treatment, differences in the beat rate of the hiPSC-CMs between the mono- and triple culture conditions were also observed (Fig. 1D). Contraction traces were analyzed using CardioMotion software 24 h after the first and second treatments with either 1 or 3 μM Doxo [15]. Contractility was not assessed after the third cycle of Doxo treatment due to significant cell death. Although contraction amplitudes did not initially differ significantly between culture types, co-cultures had notably lower amplitudes after the second Doxo treatment (Fig. 1E). Additionally, analysis of contraction duration parameters, specifically relaxation time and time-to-peak (Figs. 1F and 1G), revealed significantly shorter relaxation times in hiPSC-CMs within the triple cultures after 1 round of Doxo treatment, suggesting more pronounced cardiotoxicity in the presence of ECs and cFBs

To visualize the effect of Doxo on the hPSC-CMs directly, we used an NKX2.5-eGFP human embryonic stem cell (hESC) reporter line in which cells differentiating to cardiomyocytes express eGFP [21]. This confirmed the increased sensitivity of triple cultures to Doxo compared to monocultures, with overall cell confluency significantly decreasing after the first Doxo treatment (Figs. 2A and 2B). The decrease in GFP area (corresponding to NKX2.5^+^ hESC-CMs) paralleled the decrease in phase confluency, indicating that the hESC-CMs were also dying sooner in response to Doxo when co-cultured with the other cardiac cell types (Fig. 2C). However, again here it was challenging to accurately quantify cell death with conventional phase microscopy analysis. For example, in cultures exposed to multiple treatment rounds of 1 μM Doxo, the monocultures appeared to proliferate due to increased cell spreading before a decrease in confluency after 80 h was observed (Figs. 2B and 2C). Furthermore, while visual inspection confirmed 100% cell death at the last timepoint, quantitative analysis based on confluency only indicated a ∼60% reduction in both phase confluency and GFP area.

Because of these limitations, we investigated whether we could quantify caspase activation as a measure of Doxo-mediated apoptosis using a fluorescence assay. Although this was possible when hPSC-CMs were treated with a single high dose (10 μM) of Doxo (Supplementary Figs. 2B and 2C), the utility of the assay was limited in the cumulative treatment protocol with abrupt peaks in the fluorescence signal observed in all analyses performed (Supplementary Figs. 2D-G). This suboptimal fluorescence quantification was likely due to artefacts introduced with the re-addition of the dye after each medium replacement and the loss of labelled cells with washes.

Our findings thus indicated that hPSC-CMs in multi-cell type cultures of cFbs and ECs were more sensitive to Doxo-induced toxicity than hPSC-CMs cultured alone. However, due to the complexity of the treatment protocol, accurate quantification was not possible using standard image processing assays indicating the need for alternative analytical techniques, which we next sought to address.

### *In silico* prediction software can accurately detect and quantify hPSC-CMs and caspase activation

We investigated whether the deep neutral network (DNN) tools could facilitate and improve the quantification of toxicity evident in the phase images, as well as identify specific cell types. When comparing fluorescence images of multi-cell type cultures that contained NKX2.5-eGFP^+^ hESC-CMs, hiPSC-cFBs and - ECs, and exposed to different treatments, with images in which the NKX2.5-expressing cells were determined using the DNN, we observed a mean accuracy of 74.4% across all predictions (Figs. 3A and 3B). Further analysis confirmed that *in silico* NKX2.5 predictions closely matched actual NKX2.5-eGFP measurements in multi-cell type cultures treated with either a lethal concentration of Doxo (10 μM) or DMSO (Fig. 3C).

**Figure 3.**
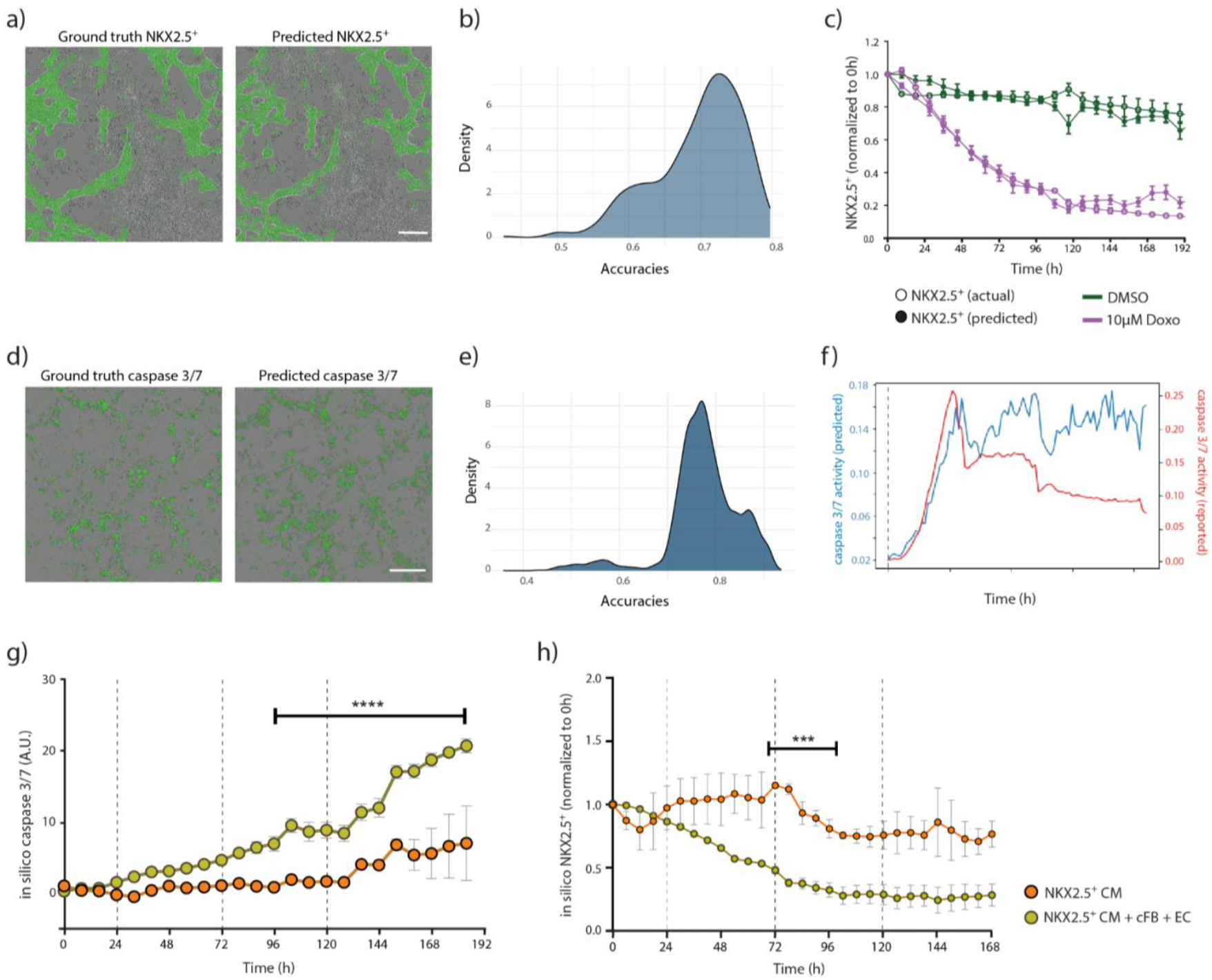
Validating the DNN tools for cardiomyocyte and caspase activity prediction. **a)** Matched images comparing actual NKX2.5-eGFP^+^ hESC-CMs (ground truth) merged with phase contrast to the corresponding *in silico* predicted NKX2.5-eGFP^+^ cardiomyocytes derived from phase contrast imaging. Scale bar, 200 μm. **b)** Analysis evaluating the distribution of the accuracy of predicted in silico masks against a total of 7200 actual NKX2.5-eGFP^+^ images. **c)** Graph depicting the overlap between *in silico* NKX2.5^+^ predictions and actual NKX2.5^+^ measurements over time, normalized to baseline (0h). Lines represent control cells treated with DMSO (green) and those exposed to single 10 μM Doxo treatment (purple). **d)** Matched images comparing actual caspase 3/7 expression (ground truth) merged with phase contrast to the corresponding *in silico* predicted caspase 3/7 expression derived from phase contrast imaging. Scale bar, 200 μm. **e)** Analysis evaluating the distribution of the accuracy of predicted in silico masks against known caspase 3/7 fluorescent dye values from a total of 8537 images. **f)** Time-course comparison of caspase 3/7 activity, contrasting *in silico* predictions with quantification obtained for caspase 3/7 dye-labelled cells treated with 10 µM Doxo. **g-h)** *In silico* quantification of caspase 3/7 activity **(g)** and NKX2.5-eGFP^+^ hESC-CMs, normalized to baseline (0h) **(h)**, in monoculture versus multi-cell type culture conditions undergoing cumulative treatment with 1 µM Doxo (dotted lines). Asterisks and black bars indicate the time points where there are statistically significant differences between the monoculture and the multi-cell type culture setups. Statistical analysis was performed with two-way ANOVA and Sidak’s multiple comparisons for each time point. Analysis is based on 3 biological replicates, each with 3 technical replicates, with error bars representing SEM.

Additionally, we applied a DNN to identify cells undergoing caspase 3/7-mediated apoptosis from phase contrast images. Also here, the hPSC-derived cells that were predicted by the DNN analysis to express caspase 3/7 closely matched the actual labelling, with a mean accuracy of 77.3% across the various cell types, time points and treatments analyzed (Figs. 3D and 3E). Quantification using the DNN also appeared more reliable, detecting high levels of caspase activity over the entire duration of the experiment, compared to the caspase 3/7 dye in which the signal intensity waned over time, particularly after medium changes (Fig. 3F).

We then reanalyzed the phase contrast and fluorescence imaging data collected of the mono- and triple-cultures of the NKX2.5^eGFP^ hESC-CMs treated with multiple rounds of 1 μM Doxo (Fig. 2) using the DNN tools. Quantification better reflected the visual observation; namely that triple cultures were more sensitive to Doxo-mediated apoptosis (Fig. 3G), and that the hPSC-CM numbers declined more quickly in these wells than in monocultures (Fig. 3H). We also analyzed data collected from hPSC-CMs exposed to both toxic and non-toxic cardiac relevant drugs using the DNN caspase 3/7 software (Supplementary Fig. 3). As expected, caspase-3/7 activity was only predicted in the cultures treated with either Doxo or ouabain, a Na^+^/K^+^-ATPase inhibitor also known for its cardiotoxicity [22]. Other compounds, such as isoprenaline and nifedipine as well as DMSO, were not predicted to induce caspase-3/7 activity, consistent with observations in cultures labelled with the caspase-3/7 dye (Supplementary Fig. 3A).

### Differential Doxo toxicity responses between isogenic cardiac and non-cardiac cell types

To investigate whether the higher sensitivity of cells in the triple cultures to Doxo might be due to differential sensitivity of one cell type, we separately exposed each cell type to the cumulative Doxo treatment. We also included hiPSC-derived dermal fibroblasts (dFBs), differentiated from the same hiPSC line, as a non-cardiac cell type for comparison. Phase contrast images clearly indicated that ECs were affected after a single round of exposure to 1 µM Doxo, while cFBs showed delayed cumulative effects (Fig. 4A). Interestingly, the hiPSC-dFBs were not only resistant to the cumulative Doxo treatment but also continued to proliferate.

**Fig 4.**
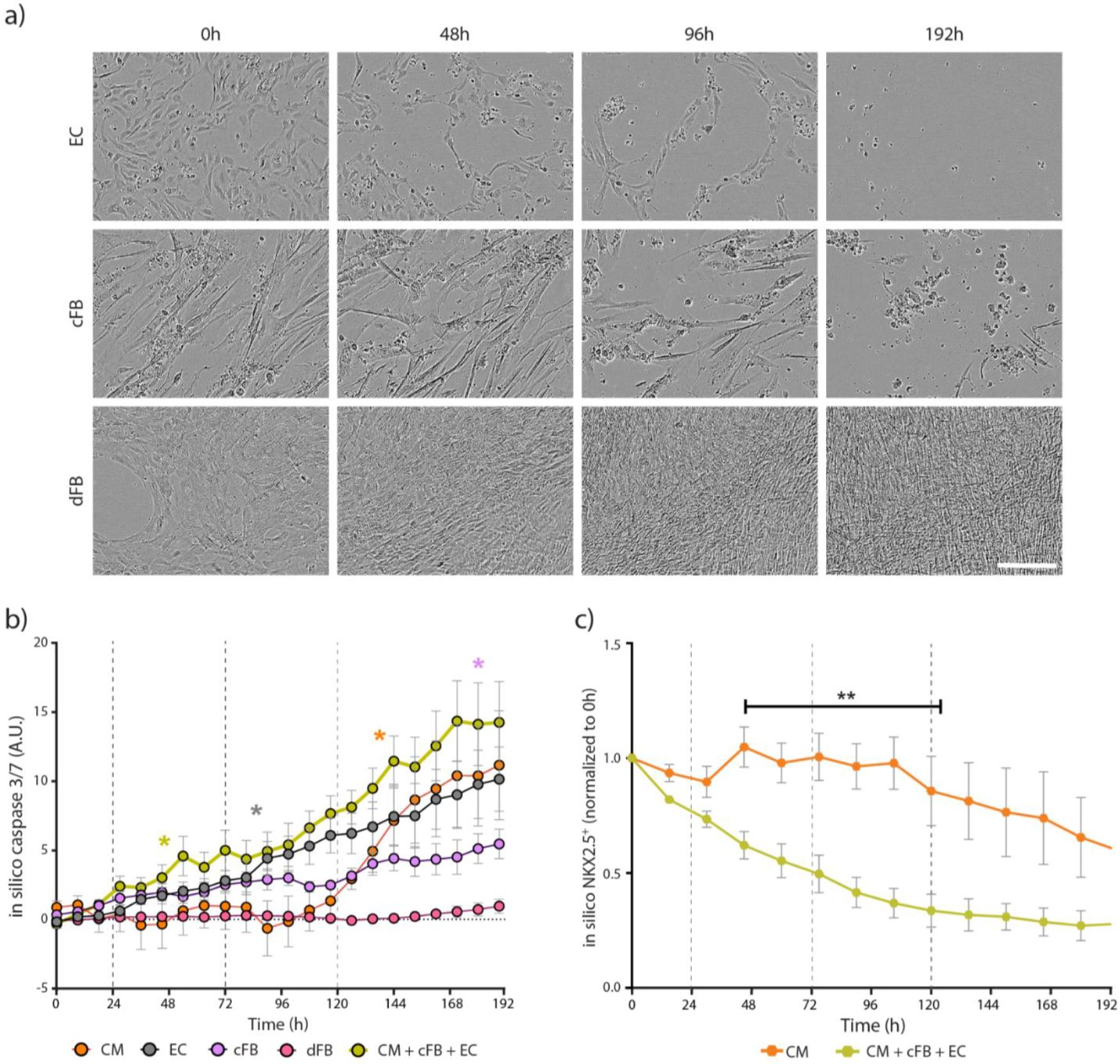
Differential Doxo sensitivity of individual isogenic hiPSC-derived cell types. **a)** Representative phase contrast images of monocultures of hiPSC-ECs (top panel), -cFBs (middle panel), and -dFBs (bottom panel) undergoing cumulative treatment with 1 µM Doxo. Images were acquired at time points corresponding to baseline (0h), 24 h after Treatment 1 and 2 (48h and 96h, respectively), and the final time point (192h). Scale bar, 200 μm. **b)** *In silico* quantification of caspase 3/7 activity for all isogenic monocultures, as well as the tri-cellular culture condition, undergoing cumulative treatment with 1 µM Doxo (dotted lines). Color-coded asterisks indicate the initial time point at which caspase 3/7 activity is significantly higher than baseline (0h) for each cell type. **c)** *In silico* quantification of NKX2.5^+^ cells in monoculture hiPSC-CMs versus tri-cell type culture conditions. The asterisk and black bar indicate the time points where there are statistically significant differences between the 2 cultures. Statistical analysis was performed with two-way ANOVA analysis without multiple comparisons or with Sidak’s corrections for multiple comparisons. Analysis is based on 3 biological replicates, each with 3 technical replicates, with error bars representing SEM.

Quantitative evaluation using the DNN caspase 3/7 tool supported the above observations, as well as the differential response observed in Fig. 1B with the hiPSC-CM monocultures and triple cultures from this cell line (Fig. 4B). Distinct temporal toxicity profiles were detected for each cell type, indicating variable sensitivities to Doxo. Among the individual cell types, hiPSC-ECs were the most sensitive, displaying significant *in silico* caspase 3/7 activity within 75 h of the initial Doxo treatment. In contrast, both cFBs and dFBs showed overall less caspase activation, with dFBs expressing almost no caspase 3/7, reflecting observations in the phase contrast images. To determine if the toxicity resistance of the hiPSC-dFBs was Doxo specific, we treated these cells with Carfilzomib, a proteasome inhibitor used clinically to treat multiple myeloma but also known to cause broad tissue toxicity [23]. In contrast to their Doxo response, dFBs were sensitive to Carfilzomib, showing significant cell death and exhibiting high levels of *in silico* caspase 3/7 activity within 24 h of treatment (Supplementary Figs. 4A-4C). Additionally, the caspase activity of triple cultures in which the hiPSC-cFBs were substituted for hiPSC-dFBs were compared (Supplementary Fig. 4D). DNN caspase 3/7 analysis indicated no significant differences in caspase levels between the triple cultures.

As previously seen with the NKX2.5-eGFP^+^ hESC-CMs (Fig. 3G), caspase 3/7 activity was detected in the triple culture following the first treatment, but only became apparent in the hiPSC-CM monoculture after the third round of Doxo exposure (Fig. 4B). Further, *in silico* prediction of NKX2.5 confirmed that apoptosis in hiPSC-CMs occurred significantly faster when these cells were co-cultured with other cell types (Fig. 4C). Detailed Doxo dose-response and time course analysis for *in silico* caspase 3/7 activity, confirmed that repeated exposures to Doxo at concentrations below 1 µM resulted in minimal effects after 3 cycles across most conditions tested (Supplementary Figs. 5A-5C). An exception was monocultures of hiPSC-ECs, with noticeable caspase activity detected after 3 rounds of 0.03 µM Doxo treatment (Supplementary Fig. 5C). At the higher concentration of 3 µM, Doxo induced significant caspase 3/7 activity from just 1 treatment cycle in hiPSC-CM and -EC monocultures as well as in the multi-cell type culture, while hiPSC-cFBs required 2 cycles (Supplementary Fig. 5D). Treatment with 10 μM Doxo quickly resulted in significant caspase 3/7 activity in hiPSC-EC and multi-cell type cultures within 6 h of treatment, and within 2 days of the single treatment in monocultures of hiPSC-CMs and -cFBs (Supplementary Fig. 5E). In contrast, hiPSC-dFBs did not show consistent and significant caspase 3/7 activity at any Doxo concentration.

### hiPSC-ECs amplify doxorubicin-induced cardiotoxicity in co-culture systems

Given the greater sensitivity of the hiPSC-ECs to Doxo, we hypothesized that these cells contributed to the increased caspase 3/7 activity observed in the multi-cell type cultures. To test this, we treated dual-cell cultures of hiPSC-CMs with either ECs or cFBs to either a single exposure of 10 μM Doxo or the cumulative 1 μM treatment protocol. Phase contrast images clearly showed that within 48 h of the initial Doxo treatment, many cells in the hiPSC-CM and -EC co-cultures appeared apoptotic (Fig. 5A). While cell death was also observed in the co-culture of hiPSC-CMs and -cFBs, it was less pronounced and only observed with the 10 μM Doxo treatment. This was quantitatively supported by the DNN caspase 3/7 analysis, which showed a significantly earlier caspase response in the cultures containing hiPSC-ECs following treatment with 10 μM Doxo, and an overall significantly higher level of caspase activity in those exposed to the cumulative 1 μM Doxo treatment (Figs. 5B and 5C).

**Fig 5.**
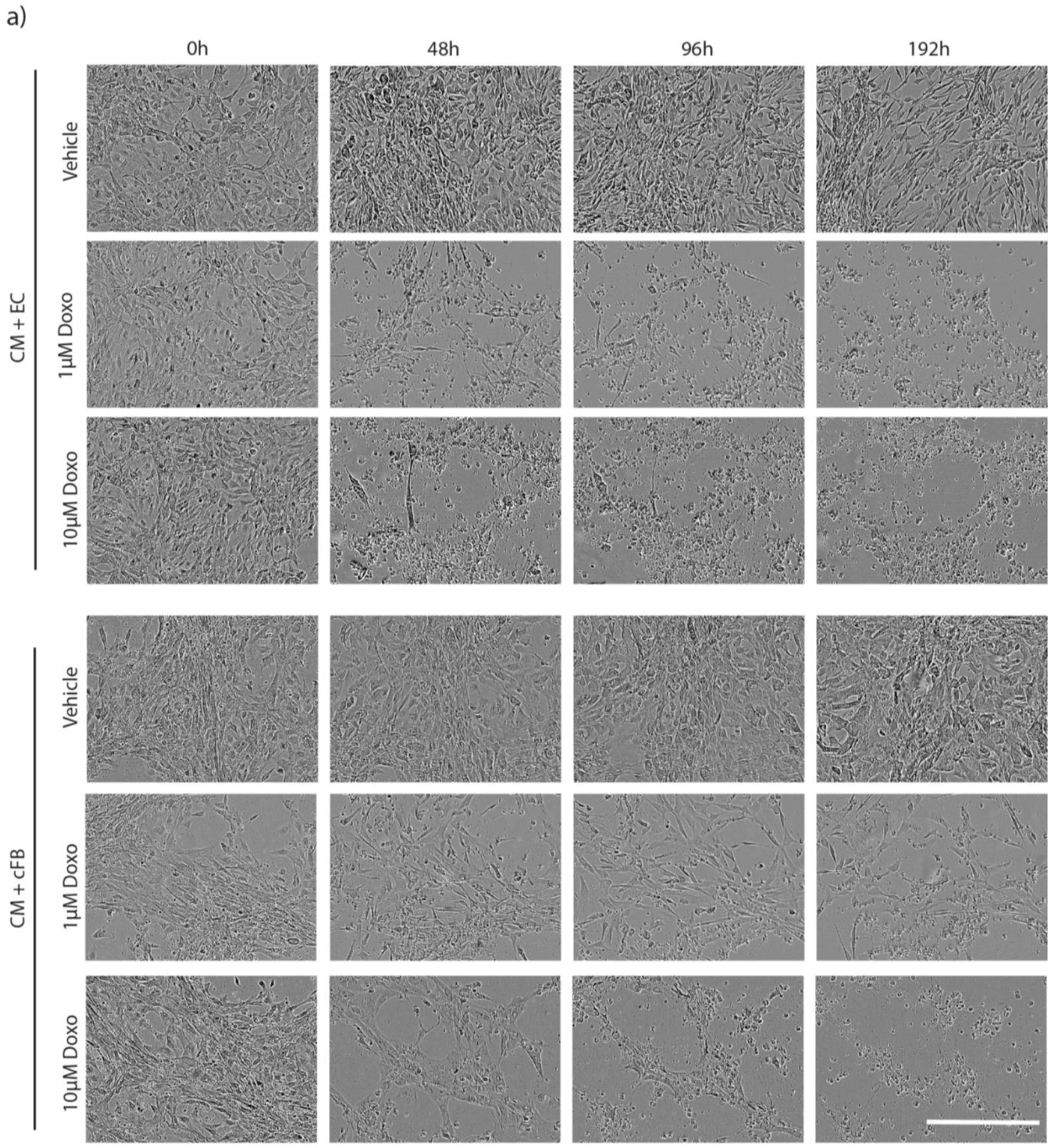
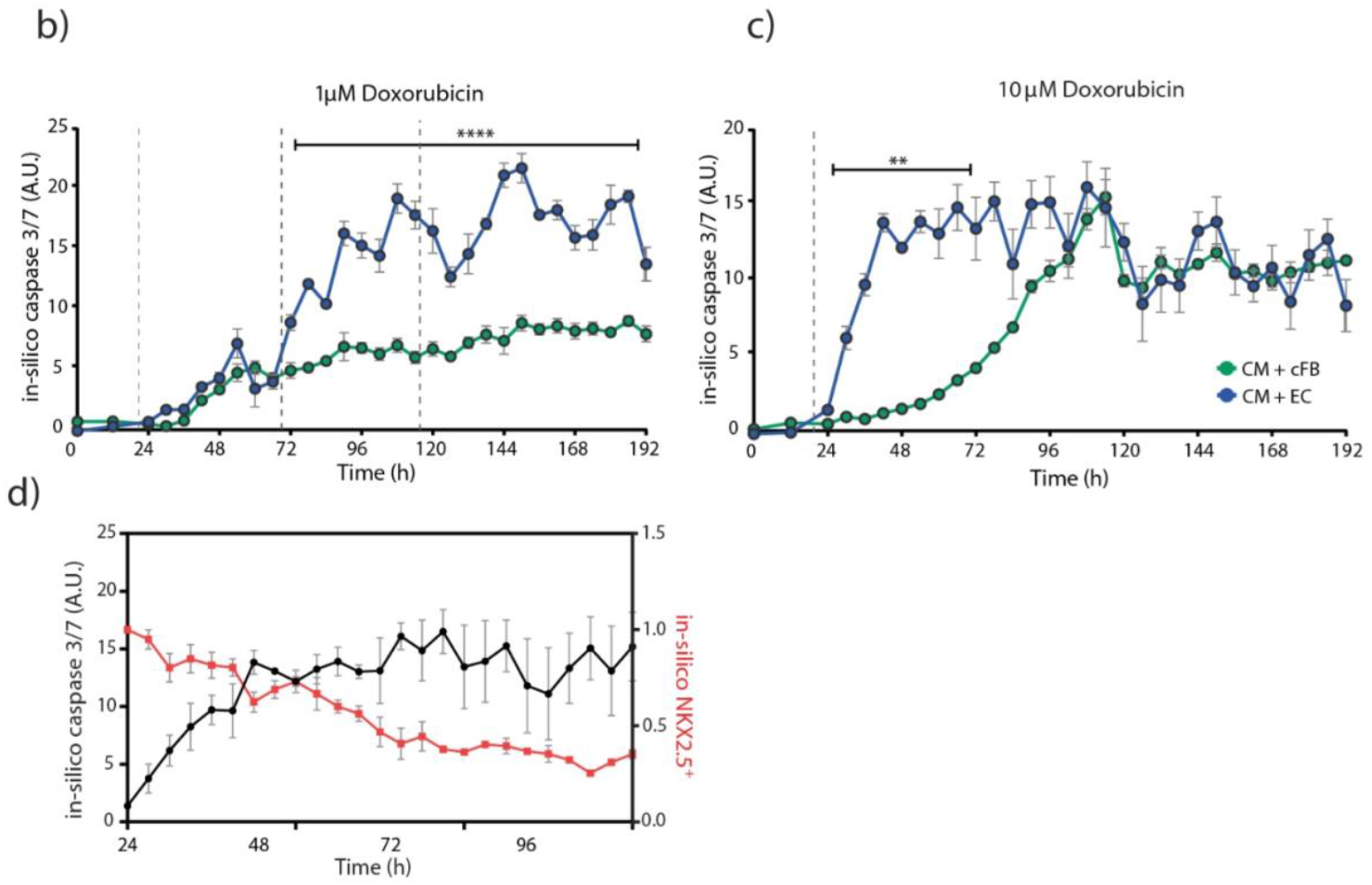
Contribution of hiPSC-ECs and -cFBs to Doxo-induced cardiotoxicity. **a)** Representative phase contrast images of hiPSC-CMs co-cultured with isogenic hiPSC-ECs or hiPSC-cFBs treated with 1 µM or 10 µM Doxo. Images were acquired at time points corresponding to baseline (0h), 24 h after Treatment 1 and 2 (48h and 96h, respectively), and the final time point (192h). 10 µM Doxo was only administered once. Scale bar, 200 μm. **b-c)** *In silico* quantification of caspase 3/7 activity in either co-cultures of hiPSC-CMs with hiPSC-ECs or -cFbs and treated with either 1 µM **(b)** or 10 µM **(c)** Doxo. Dotted lines indicate the treatment time points, while the asterisks and black bars indicate the time points where there are statistically significant differences between the 2 co-cultures. **d)** Simultaneous *in silico* quantification of caspase 3/7 activity (black line) and NKX2.5^+^ cells (red line) in co-cultures of hiPSC-CMs and -ECs treated with 1 µM Doxo at 24 and 72 h time points, comparing the dynamics of apoptosis with rate of cardiomyocyte death. Statistical analysis was performed with two-way ANOVA and Sidak’s multiple comparisons. Analysis is based on 3 biological replicates, each with 1 or 2 technical replicates, with error bars representing SEM.

Using the DNN tools, we also examined the specific effect on the hiPSC-CMs within the co-cultures treated with 1 µM Doxo (Fig. 5D). As previously observed, there was a sharp increase in caspase 3/7 activity (and general cell death) within the first 24 h of treatment that plateaued thereafter. In contrast, hiPSC-CMs levels steadily decreased throughout the whole treatment, suggesting that Doxo has an acute impact on the hiPSC-ECs but gradually affects the hiPSC-CMs. Overall, these results indicated interplay between the hiPSC-CMs and hiPSC-ECs in mediating Doxo-induced cardiotoxicity and suggests that ECs may have a pivotal role in exacerbating cardiomyocyte sensitivity to Doxo.

## Discussion

Despite its effectiveness in a broad range of cancers, Doxo is notorious for its cardiotoxic side effects which can manifest in some patients shortly after treatment begins or as late as 20 years after therapy completion [24]. While the cumulative dose has been identified as a critical predictor of Doxo-induced cardiac dysfunction, clinical attempts to limit the total dose have primarily succeeded in reducing instances of acute toxicity without diminishing the incidence of late onset effects [25]. The challenge of mitigating these effects, while maintaining therapeutic efficacy, underscores a pressing need for more accurate models to understand Doxo-induced cardiotoxicity and that of its analogues [26].

Combining hiPSC-derived cardiac cells with appropriate treatment protocols, is a promising approach to bridge the gap between *in vitro* studies and clinical reality. In particular hiPSC-cardiac cells could be a powerful model for understanding variability in the cumulative dose tolerances of patients to Doxo, which in part is attributed to genetic differences [27]. Here we monitored the cumulative effect of Doxo over time by incubating the cells for 4 h with 1 µM Doxo every 48 h over the course of 8 days. This scheme was based on the reported pharmacokinetics of the drug *in vivo*, where the free-plasma concentration of Doxo rapidly drops from initial levels ranging between 20 nM to 2 µM, and has a terminal half-life of 20-48 h [18-20].

3D cardiac models incorporating hiPSC-CMs, such as microtissues or engineered heart tissues, are increasingly being used to evaluate cardiotoxicity *in vitro* [2,17,28,29]. These models not only enhance cardiomyocyte maturation but also facilitate intercellular crosstalk between key cardiac cell types, which is crucial not only for cardiac function but also for examining toxicity responses [25,30-33]. Studies using these 3D models have provided valuable insights, particularly regarding functional analysis. For example, tissues exposed to Doxo demonstrated compromised functionality within 24 h of treatment [2,17]. Conversely, alternative drugs such as aclarubicin, formulated to be less cardiotoxic, only affected contractility and force of contraction at later time points.

While these models hold promise, several challenges still hinder their utility as drug screening tools. These include scalability issues, ensuring uniform accessibility of the compounds to all cells, and difficulties in obtaining comprehensive, cell type-specific responses to the effects of the compound. This led us to develop a 2D model that still captures the multicellular dynamics of the heart by co-culturing various hPSC-derived cardiac cells, but also allows easier imaging and monitoring. This approach indicated that cardiomyocytes alone, conventionally the focus in cardiotoxicity studies, are not the most sensitive cardiac cell type affected by Doxo, requiring 3 treatment rounds for visible cell death. Instead, when cultured together with hiPSC-cFBs and -ECs, signs of apoptosis based on caspase 3/7 activity appeared within a few hours after the initial Doxo exposure, suggesting the presence of other cell types exacerbated hPSC-CM toxicity. This was also reflected by the more rapid changes in their contractility observed in these multi-cell type conditions, underscoring the intricate interplay between different cardiac cell types in mediating the toxic effects of Doxo.

Our findings align with previous reports indicating that cFBs and ECs exhibit early signs of Doxo-induced stress. These studies identified a pro-fibrotic influence of Doxo on fibroblasts in rat hearts via TGF-β and SMAD3 signaling, triggering their transition to myofibroblasts [34]. Furthermore, mouse cFBs exposed to Doxo were reported to become senescent leading to the secretion of pro-inflammatory cytokines, such as IL-1β [35]. Likewise, clinical studies have associated Doxo cardiotoxicity with damage to the vascular endothelium [36,37], with *in vitro* or animal studies indicating Doxo can compromise endothelial elasticity, increase cardiac microvasculature permeability, inhibit vascular network formation, and induce oxidative stress through altered reactive oxygen species and nitric oxide (NO) levels [25].

Interestingly, Doxo-mediated toxicity occurred earlier when hPSC-CMs were co-cultured with hiPSC-ECs compared to co-cultures of hPSC-CMs with -cFBs. Recently, we confirmed crosstalk between hiPSC-ECs and -CMs via the NO-pathway enhanced inflammatory responses and influenced the contractility of the hiPSC-CMs [38]. It is possible that disrupted communication between these cells triggered by the exposure to Doxo alters paracrine signaling, particularly involving NO, and this could be responsible for the enhanced cardiomyocyte toxicity observed when cultured together with the hiPSC-ECs.

Live-cell imaging often employs organic fluorophores or fluorescent proteins to monitor specific cellular processes or markers [39]. However, a common limitation, which we also encountered with the caspase 3/7 dye, is the diminishing fluorescence signal over time, particularly following culture medium changes. This becomes increasingly significant during lengthy treatment protocols involving multiple drug administrations, as demonstrated in this study. While tagging endogenous markers with fluorescent proteins can mitigate some of these concerns, such assays are generally restricted to the modified cell lines in which they are developed, thereby limiting the ability to assess patient-specific responses.

To overcome these issues, we applied a DNN designed for apoptosis analysis to quantify *in silico* caspase 3/7 activity from phase-contrast images. This allowed us to circumvent the limitations of the dye-based assays, enabling continuous monitoring of drug-induced toxicity over extended periods and the analysis of effects across different cell types and within multi-cell type combinations without the need for direct labeling. Moreover, the ability of the tool to segment and classify cells accurately from phase contrast images facilitated the identification of hPSC-CMs within mixed cultures of cardiac cells. This was invaluable for not only tracking cardiomyocyte toxicity from different hPSC lines over time, but also revealing the varying degrees of toxicity when the hPSC-CMs were co-cultured with different cell types.

The use of ML for assessing cardiotoxicity, particularly in hPSC-CMs, is advancing rapidly. For example, ML models have demonstrated high accuracy in distinguishing cardiotoxic and non-cardiotoxic compounds based on functional parameters like action potential [40], calcium cycling [41] and contraction properties [42,43]. Additionally, deep learning models have been developed to assess cardiotoxicity from high-content imaging data of hiPSC-CMs treated with a range of pharmacological compounds, combined with contractility data [42,44]. However, in these instances the hiPSC-CMs are fixed, with sarcomere organization and alignment evaluated by immunofluorescence. Nevertheless, Maddah *et al*. was able to demonstrate that with continuous exposure to low concentrations of Doxo (0.3 – 0.6 μM), the resulting toxicity could be detected in phase-contrast images of live hiPSC-CMs with greater sensitivity than cell confluency assays [42]. However, it is unclear whether their image-based deep-learning software would be able to accurately detect toxicity in treatment strategies that involve repeated short exposures to compounds along with media changes, such as performed here.

Indeed, applying ML in biological contexts demands careful feature selection, model training, and accurate cell segmentation, which can be challenging as frequently cells cluster or overlap [45]. Nonetheless, several recent studies have successfully used ML to identify and classify different cell types, such as macrophage subtypes [46], as well as evaluate the suitability of hPSC-CMs for downstream experiments [47], and predict the efficiency of hPSC differentiation to cardiomyocytes [48]. Such tools, including the one we have used here to monitor hPSC-CMs within a multi-cell type culture and detect early-stage aspects of apoptosis from phase contrast images, substantially streamlines experimental design and enhances the efficiency of cardiotoxicity evaluations.

In summary, while hPSC-CMs have long been the focus of cardiotoxicity studies, our research suggests that the inclusion of other cardiac cell types, in particular hiPSC-ECs, critically influences the cardiotoxic outcome of Doxo treatment. These findings, enabled by the application of ML in 2D cell culture models, not only indicates that cells other than cardiomyocytes play a role but also suggests new cellular targets for preventing Doxo-induced cardiotoxicity.

