## Supplementary documents for "A simplified co-culture reveals altered cardiotoxic responses to doxorubicin in hPSC-derived cardiomyocytes in the presence of endothelial cells"

**Supplementary Figure 1. Effect of cumulative doxorubicin treatment on hiPSC-CMs**

**Supplementary Figure 2. Difficulties in accurately quantifying confluency and caspase 3/7 activity in hiPSC cardiac cultures**

**Supplementary Figure 3. Analysis of caspase 3/7 activity in hiPSC-CMs following treatment with various compounds**

**Supplementary Figure 4. Evaluation of toxicity in cultures containing hiPSC-dermal fibroblasts**

**Supplementary Figure 5. *In silico* quantification of caspase 3/7 activity in isogenic hiPSC-derived cells**

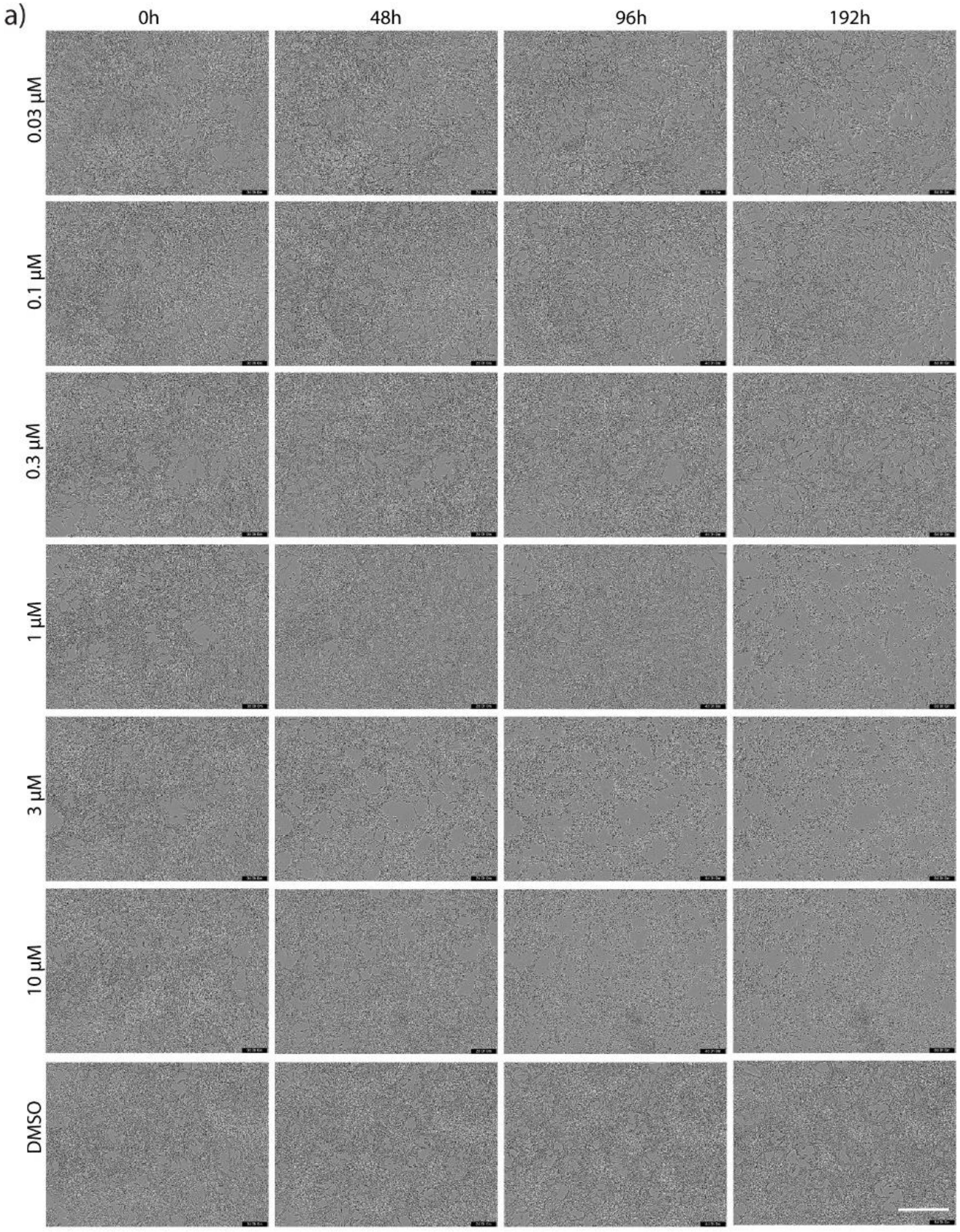
a)
0h
48h
96h
192h
0.03 μM
0.1 μM
0.3 μM
1 μM
3 μM
10 μM
DMSO

b)

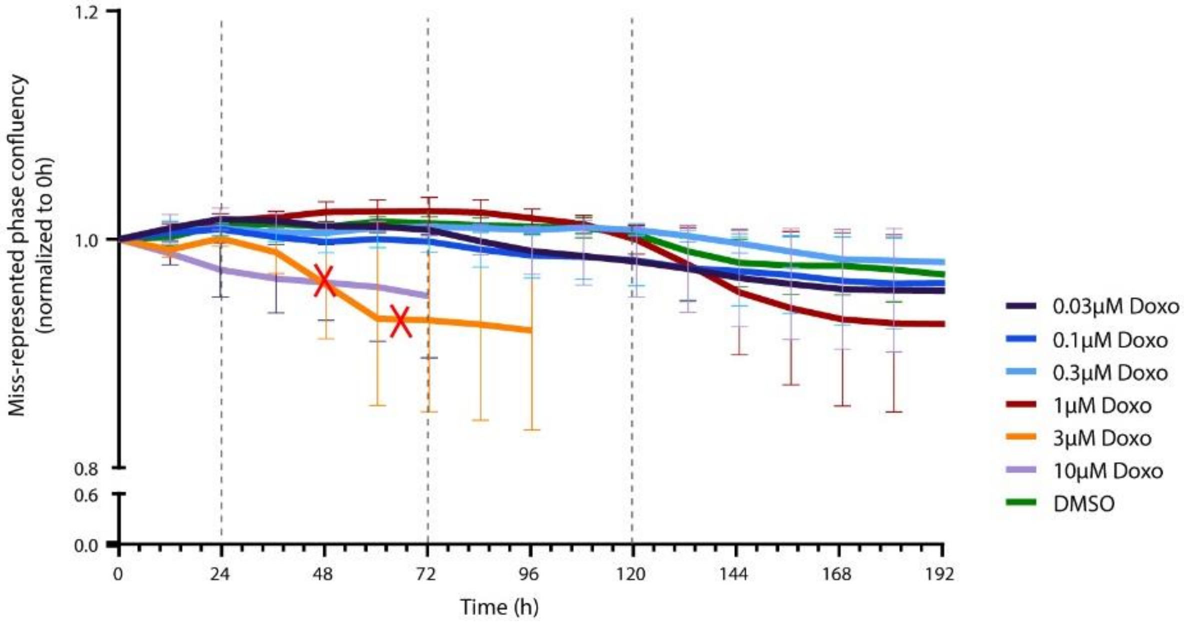


**Supplementary Fig 1. Effect of cumulative Doxo treatment on hiPSC-CMs.**

**a)** Representative phase contrast images of monoculture hiPSC-CM cumulatively treated with various Doxo concentrations or DMSO (vehicle control). Images were acquired at time points corresponding to baseline (0h), 24 h after Treatment 1 and 2 (48h and 96h, respectively), and the final time point (192h). All treatments were performed according to the cumulative treatment protocol outlined in Fig. 1, except 10µM Doxo, which was added once. Scale bar, 400µm. **b)** Incucyte quantification of confluence (percentage of the image area covered by objects) was normalized to baseline (0 h). Red crosses indicate the time points at which all cells were visually observed to be dead. Dotted lines indicate treatment time points. Analysis is based on 3 biological replicates, each with 3 technical replicates, with error bars representing SEM.

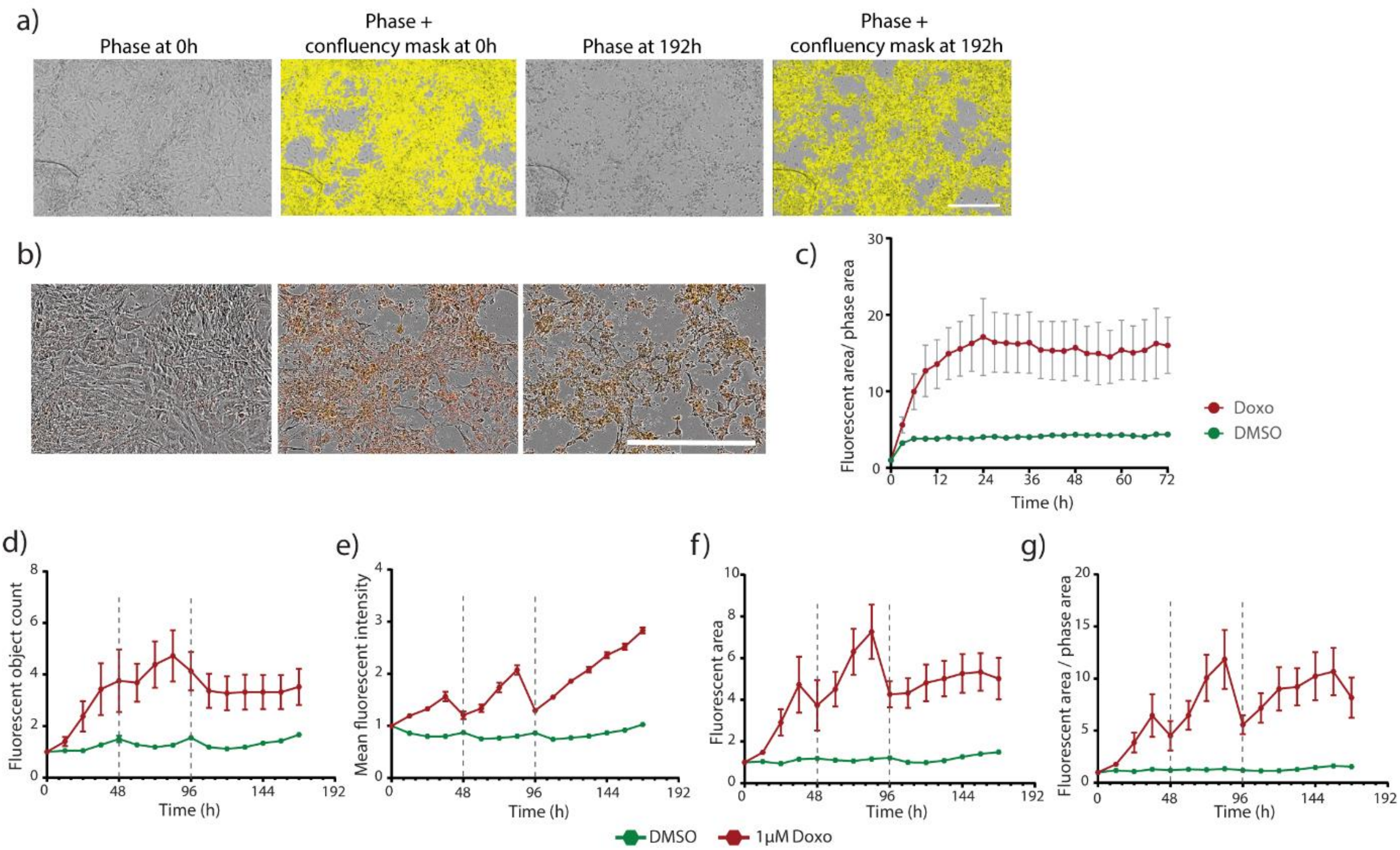


**Supplementary Fig 2. Difficulties in accurately quantifying confluency and caspase 3/7 activity in hiPSC cardiac cultures.**

**a)** Representative phase contrast images of multi-cell type cultures (hiPSC-CMs, -ECs, and -cFBs), with the corresponding phase confluency masks (highlighted in yellow) generated following analysis highlighting the inability of confluency masks to distinguish live and dead cells. Images are shown for at the start (baseline, 0h) and at the final time point (192h) following 1 μM cumulative Doxo treatment, at which point all cells visually were deemed to be non-viable. Scale bar, 400 μm. **b)** Merged phase and red fluorescence images of hiPSC-CMs labelled with caspase 3/7 dye and treated once with 10 μM Doxo. Time points correspond to baseline (0h), 24 h and 72 h after treatment. Scale bar, 200 μm. **c)** Time-course quantification of caspase 3/7 fluorescence area, relative to the phase contrast area quantified from the confluency mask, for hiPSC-CMs treated with 10 μM Doxo or DMSO (vehicle control). **d-g)** Evaluation of various metrics for quantifying the temporal changes in caspase 3/7, based on fluorescence, in multi-cell type culture conditions undergoing cumulative treatment with 1 μM Doxo or DMSO (vehicle control). Dotted lines indicate the treatment time points. In all analyses measurements were normalized to baseline (0 h). Analysis is based on 3 technical replicates, with error bars representing SEM.

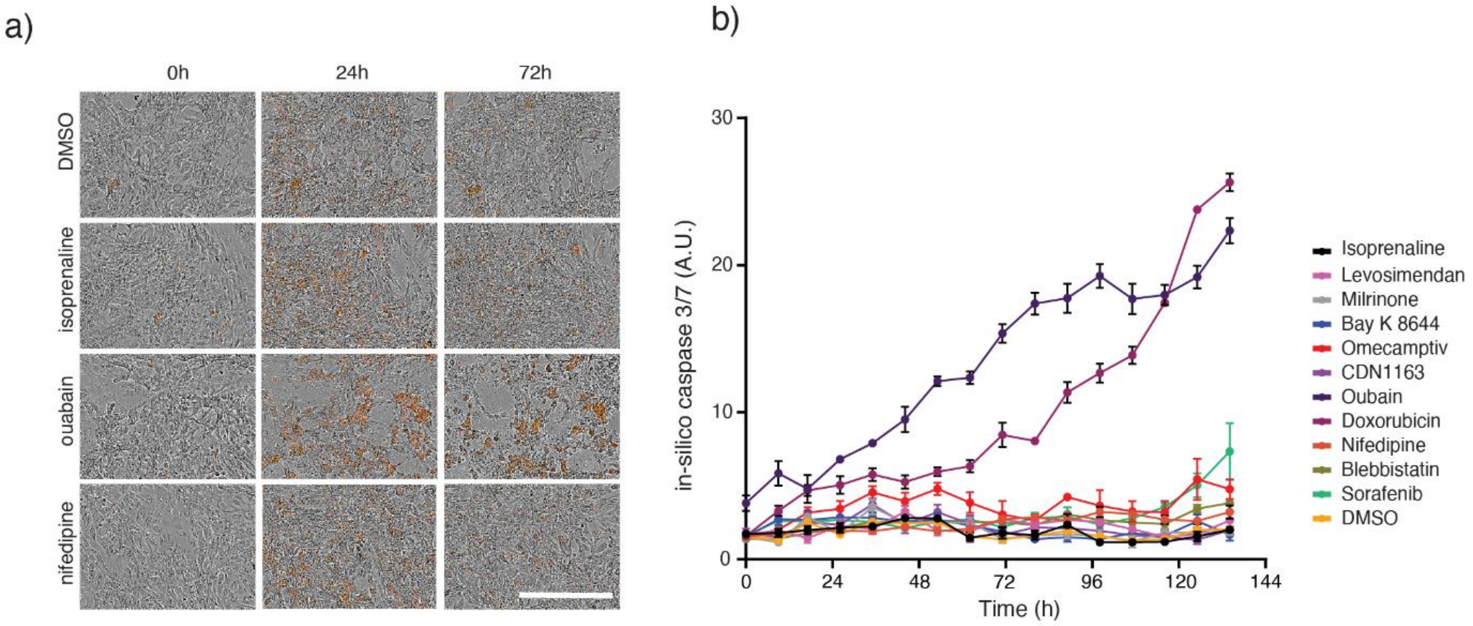


**Supplementary Fig 3. Analysis of caspase 3/7 activity in hiPSC-CMs following treatment with various compounds.**

**a)** Merged phase and red fluorescence images of hiPSC-CMs labelled with caspase 3/7 dye and treated once with the indicated compounds. Time points correspond to baseline (0h), 24 h and 72 h after treatment. scale bar, 400 µm. **b)** *In silico* quantification of caspase 3/7 activity over time in hiPSC-CMs treated once with indicated compounds. The treatment concentrations for the compounds are listed in Supplementary Table 1. Analysis is based on 3 biological replicates, with error bars representing SEM.

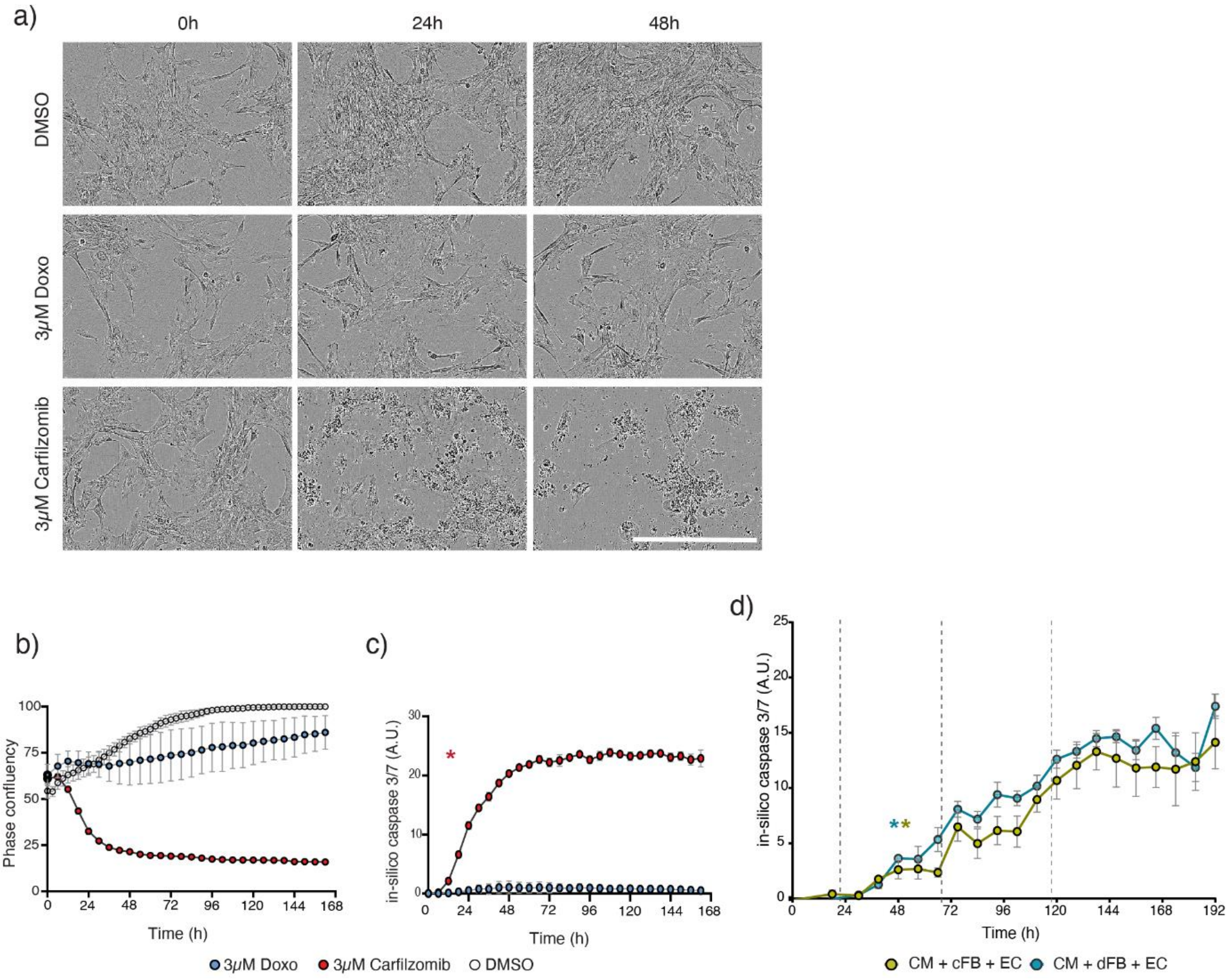


**Supplementary Fig 4. Evaluation of toxicity in cultures containing hiPSC-dFbs.**

**a)** Representative phase contrast images of hiPSC-dFbs treated with either vehicle control (3 µM DMSO), Doxo or Carfilzomib. Images were acquired at time points corresponding to baseline (0h), and 24 h and 48 h after treatment. Scale bar, 400 µm. **b)** Time course quantification of phase confluency area calculated from live cell imaging for the treatment conditions described in **(a)**. **c)** *In silico* quantification of caspase 3/7 activity in hiPSC-dFbs for the treatment conditions described in **(a)**, normalized to the vehicle control. The asterisk indicates the initial time point at which caspase 3/7 activity is significantly higher than baseline (0 h). **d)** *In silico* quantification of caspase 3/7 activity in either multi-cell type culture conditions containing either hiPSC-cFBs or -dFBs, undergoing cumulative treatment with 1 µM Doxo (dotted lines). The asterisk indicates the initial time point at which caspase 3/7 activity is significantly higher than baseline (0 h). Statistical significance was determined by two-way ANOVA analysis. Graphs present mean values with SEM as error bars. Analysis is based on 3 biological replicates.

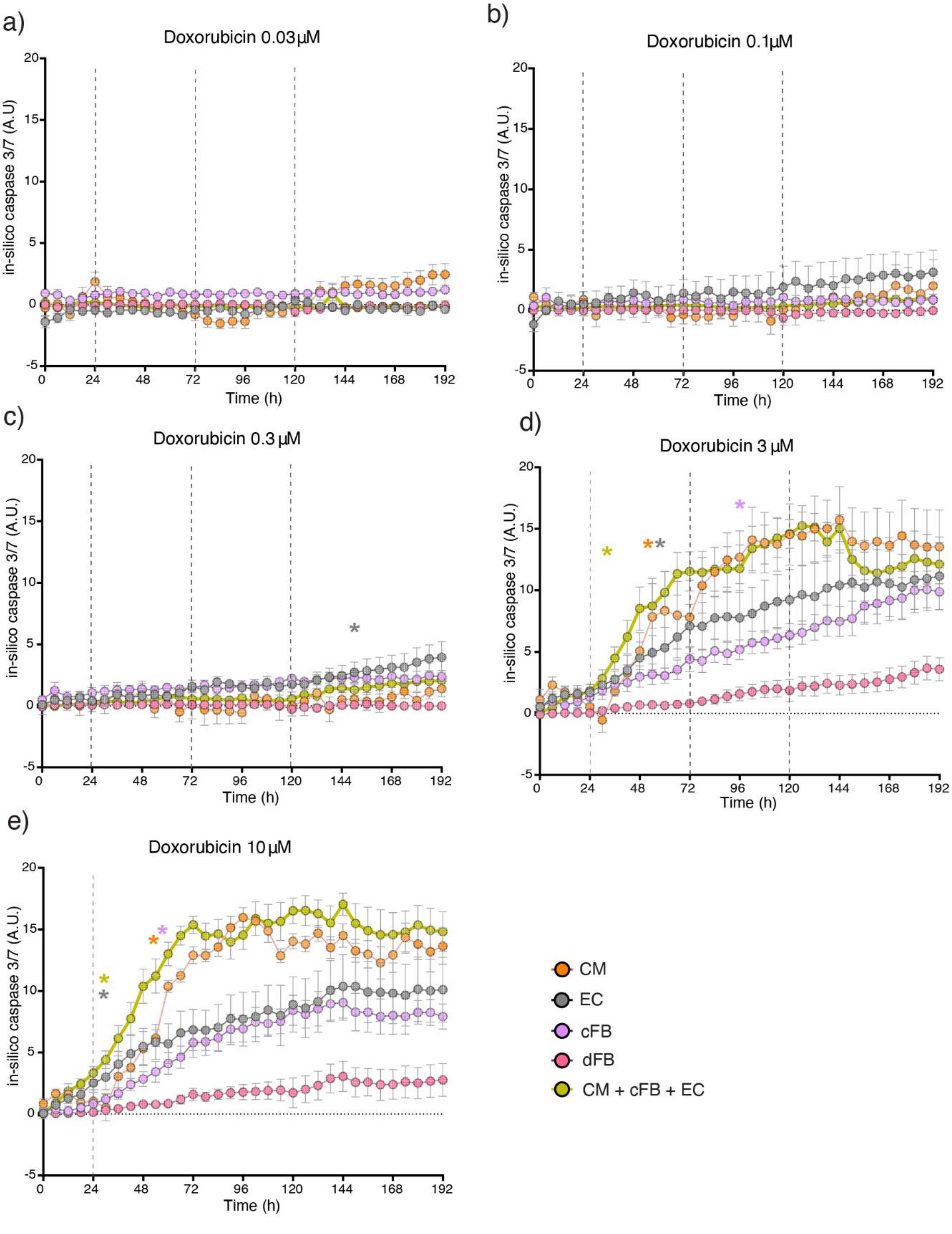
a)
Doxorubicin 0.03µM
in-silico caspase 3/7 (A.U)
Time (h)
b)
Doxorubicin 0.1µM
in-silico caspase 3/7 (A.U.)
Time (h)
c)
Doxorubicin 0.3 µM
in-silico caspase 3/7 (A.U.)
Time (h)
d)
Doxorubicin 3 µM
in-silico caspase 3/7 (A.U.)
Time (h)
e)
Doxorubicin 10 µM
in-silico caspase 3/7 (A.U.)
Time (h)
CM
EC
cFB
dFB
CM + cFB + EC

**Supplementary Fig 5. *In silico* quantification of caspase 3/7 activity in isogenic hiPSC-derived cells.** Evaluation of caspase 3/7 activity for all isogenic monocultures, as well as the multi-cell type culture condition (hiPSC-CMs, -ECs, and -cFBs) treated with different concentrations of Doxo. All treatments (dotted line) were performed according to the cumulative treatment protocol outlined in Fig. 1, except 10 µM Doxo, which was administered just once. Assessments of caspase 3/7 activity were normalized against the respective vehicle control of each culture type. Color-coded asterisks indicate the initial time point at which caspase 3/7 activity is significantly higher than baseline (0 h) for each cell type. Statistical significance was determined by two-way ANOVA analysis. Graphs present mean values with SEM as error bars. Analysis is based on 3 biological replicates, each with 3 technical replicates.
